# A highly stable engineered disulfide bond in the dimer interface of *E. coli* orotate phosphoribosyl transferase measures cytosolic redox conditions in yeast

**DOI:** 10.1101/2025.01.14.632917

**Authors:** Vilius Kurauskas, Sophia P. Albrechtsen, Sarah Gersing, Kristoffer E. Johansson, Rasmus Hartmann-Petersen, Martin Willemoës, Jakob R. Winther

## Abstract

Structural disulfides are very rarely found in cytoplasmic proteins of mesophilic organisms. Nevertheless, disulfides can form in the cytosol if these are stabilized sufficiently by the supporting protein structure. To investigate the redox properties of structural disulfide bonds, we introduced disulfide bonds into the cytosolic enzyme from *E. coli*, orotate phosphoribosyl transferase (OPRT). Because OPRT is a homodimer the introduction of opposing cysteine residues (R44C and D92C) into separate monomers of the enzyme meant that disulfide bond formation could easily be followed by non-reducing SDS-PAGE. Redox titration experiments in DTT buffers revealed that the designed disulfide bond was exceedingly stable with a redox potential of -314 mV, close to that of dithiothreitol (DTT). This interchain disulfide bond improved global thermostability by 5.9 °C relative to wild type protein and 21.4 °C above its reduced form. The catalytic activity of the enzyme in either its thiol or disulfide state was not substantially affected, indicating structural integrity. Combining an inactive subunit with the asymmetric nature of the disulfide bond enabled determining the rates of subunit rearrangement and disulfide formation. The R44C/D92C double mutant resulted in the formation of a symmetric homodimer with two interchain disulfides. However, the global stability of the two-disulfide dimer was not substantially increased relative to the dimer with the single disulfide bond, suggesting that the stabilizing effect was gained by preventing dimer dissociation. Finally, we show that the engineered disulfide bonds are formed to a significant degree in the cytosol of yeast cells, allowing us to determine the redox potential of the cytosol in equilibrium with this substrate to -300 mV. This is slightly lower than that previously determined using a GFP-based sensor, rxYFP in yeast. We find that disulfide bond formation is particularly enhanced in mutants lacking the glutathione reductase, Glr1. Thus, the engineered disulfide provides an alternative method for determining intracellular redox potential in living cells with an extended dynamic range relative to GFP-based sensors.

## Introduction

In most living organisms the formation of structural disulfide bonds is restricted to extra-cytoplasmic compartments, e.g. periplasm, endoplasmic reticulum (ER) or intermembrane space (IMS) of mitochondria (Depuydt et al., 2011). It is generally believed that the low redox potential of the cytoplasm precludes the formation of disulfides. In general, two NADPH coupled systems are employed to ensure low cytoplasmic redox potential. The first system utilizes glutathione, where a high ratio of reduced vs. oxidized state is maintained by the enzyme glutathione reductase. Glutathione reduces disulfides to form mixed protein-glutathione disulfides, which are further resolved by glutaredoxins (Grx) (Lillig et al., 2008; Jensen et al, 2014). The cytosolic redox potential of glutathione has been measured to be around -289 mV in yeast (Ostergaard et al., 2004) and -320 mV in *Arabidopsis thaliana* (Meyer et al., 2007). In the second NADPH-dependent system, substrate disulfides are reduced by thioredoxins which in turn are reduced by thioredoxin reductase (Collet and Messens, 2010).

Multiple factors may influence the high redox potential of non-native disulfide bonds, including: 1) conformational strain induced by the formation of a non-native bond (Zhou et al., 1993), 2) low effective concentration of reacting thiol groups and 3) unfavorable electrostatic factors that affect the pK_a_ of the corresponding thiols. In the folded state of a protein, however, the geometry is often optimal to achieve the highest stability of structural disulfides. Thus, the effective concentration can be extremely high, and the redox potential of structural disulfides can correspondingly be very low. Although the redox potentials are known for many oxido-reductases which employ thiol-disulfide chemistry for their activity (Ortenberg and Beckwith, 2003, Fass and Thorpe, 2018), studies on the redox properties of *native structural* disulfide bonds are exceedingly scarce in scientific literature (Burton 2000, Mitchinson and Wells, 1989).

To mimic the redox properties of a structural disulfide bond, we engineered a model disulfide at the interface of a homodimeric enzyme, orotate phosphoribosyltransferase (OPRT) from *E. coli*. Since most of the naturally occurring structural disulfides are buried (Dani et al., 2003), we wanted to create a solvent exposed disulfide bond, which would be reversibly oxidized and where reduction would not significantly disrupt structure. We here describe the successful design, and *in vitro* redox characterization of a highly stable structural disulfide whose formation is easily followed by non-reducing SDS-PAGE. We further show that a construct with two disulfide bonds provides a tool for determination of cytosolic redox conditions with an extended dynamic range.

## Materials and methods

### Chemicals

KH_2_PO_4_, MgCl_2_·6H_2_O, FeCl_3_·6H_2_O, NaH_2_PO_4_·H_2_O, HCl, Na_2_CO_3_, Na_2_B_4_O_7_·10H_2_O, ammonium persulfate and imidazole were purchased from Merck KGaA (Darmstadt, Germany). Trans-4,5-Dihydroxy-1,2-dithiane (DTT_OX_), dithiothreitol (DTT_RED_), (NH_4_)_2_SO_4_, CaCl_2_, ampicillin sodium salt, Coomassie R250, acetic acid, 5,5’-dithiobis-(2-nitrobenzoic acid), ethylenediaminetetraacetic acid (EDTA), N-cyclohexyl-3-aminopropanesulfonic acid (CAPS), 2-(N-morpholino)ethanesulfonic acid (MES), NiSO_4_·6H_2_O, orotic acid, 5-phospho-D-ribose 1-diphosphate pentasodium salt (PRPP), tris(hydroxymethyl)phosphine (THP), N-ethylmaleimide and sodium acetate were all products of Sigma-Aldrich GmbH (St. Louis, Missouri, USA). Isopropyl β-D-1-thiogalactopyranoside (IPTG), tris(hydroxymethyl)aminomethane (Tris), tetramethylethylenediamine, glycerol and acrylamide (30%, 37.5:1 acrylamide:bis-acrylamide) were products of AppliChem GmbH (Darmstadt, Germany). Yeast extract, agar, Pfu polymerase, molecular weight markers were purchased from Thermo Fisher Scientific (Waltham, Massachusetts, USA). Glucose was from Serva Electrophoresis GmbH (Heidelberg, Germany) and Bacto tryptone from Becton, Dickinson and Company (Franklin Lakes, New Jersey, USA). 2000 Da maleimide conjugated polyethylene glycol (mPEG-mal) was obtained from NOF Corporation (Tokyo, Japan).

### Cloning, expression and purification of protein variants

mutations were introduced stepwise by the QuickChange method (Stratagene, La Jolla, California, USA) to an open reading frame of wild-type *E. coli pyrE* gene with a C-terminal His_6_ tag (as described (Hansen et al., 2014)). Cys-containing variants were created by replacing original codon with TGC codon for cysteine, while native Cys residues were replaced by GCC codons for Ala. To create an inactive enzyme variant, substitutions in the flexible loop (R99G/K100S/K103S/H105G) and in the active site D124/125N were introduced in two steps. All constructed sequences were confirmed by sequencing at Eurofins MWG Operon (Ebersberg, Germany). All the primers were designed with software provided by Stratagene and listed in the supplementary information, Table S1.

Plasmids encoding mutant OPRT variants were transformed into *E. coli* strain MRH205 (Hansen et al., 2014) which is deleted for the *pyrE* gene. Cells were grown at 37 °C (after inoculation from overnight culture) in AB-LB medium (1:4 medium:flask volume) (15 mM (NH_4_)_2_SO_4_, 36.5 mM Na_2_HPO_4_, 22 mM KH_2_PO_4_, 2 mM MgCl_2_, 100 μM CaCl_2_, 10 μM FeCl_3_, 10% (w/v) tryptone, NaCl, 5% (w/v) yeast extract) supplemented with 0.2% glucose and 100 μg/ml ampicillin, until OD600 of ∼1 was reached. After addition of IPTG to 1 mM, cultures were grown for an additional 3.5 h at 37 °C and harvested by centrifugation at 5000 g for 15 minutes. Cells were resuspended in 1/10 initial culture volume of 0.9% NaCl followed by centrifugation as above. Cell pellets were stored at -20 °C. Thawed cell pellets were resuspended in 1/25 initial culture volume (20 ml) of equilibration buffer (50 mM sodium phosphate pH 8.0, 20 mM imidazole, 150 mM NaCl), sonicated on ice/water slush at 25% power for 1 minute making 2-minute breaks and repeating this 15 times using a Bandelin Electronics Sonoplus sonicator with UW 2200 ultrasonic converter and equipped with a TT 13×13 mm tip. Debris was separated by centrifugation at 12000 g for 30 minutes. Protein extract was applied to a 2 ml column of HIS-Select Nickel Affinity Gel (Sigma-Aldrich GmbH, St. Louis, Missouri, USA) equilibrated with equilibration buffer and washed with 10 ml of the same buffer. Protein was eluted with 50 mM sodium phosphate pH 8.0, 100 mM imidazole, 150 mM NaCl buffer, dialyzed twice for three hours against 50 mM Tris-HCl pH 8.0, 1 mM EDTA buffer (1:400 volume protein:buffer) at 4 °C and supplemented with 50% glycerol for storage at -20 °C.

### Determination of disulfide redox potential

Mixtures of equal amounts of OPRT ^C44*^ and OPRT^C92*^ were equilibrated in phosphate buffer pH 7.25 with varying ratios of DTT_RED_/DTT_OX_ (25-0.01) present in large excess over protein. The extent of disulfide bond formation in each buffer was evaluated by non-reducing SDS-PAGE. The formation of a disulfide bond between the OPRT^C44*^ and OPRT^C92*^ variants was well described by a simple two-state reaction:

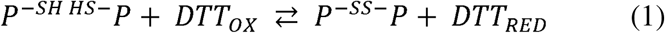

Where P^-SH^ ^SH-^P and P^-SS-^P indicates the reduced and the oxidized form of the OPRT dimer, respectively, and DTT_OX_ and DTT_RED_, the oxidized and reduced forms of DTT, respectively. Note, that even in the reduced state OPRT is a dimer and must be treated as such in mass action equations. Thus, K_OX(DTT)_ can be calculated as:

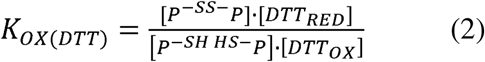

And subsequently, standard biochemical redox potentials (E°’) as determined from the Nernst equation:

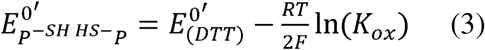

### Determination of disulfide bond formation kinetics

concentrated stocks of iOPRT^C44^ and OPRT^C92^, containing 50% glycerol, were incubated with 2 mM tris(hydroxymethyl)phosphine (THP) and 2 mM reduced DTT for 30 minutes and reductants removed on a NAP5 column (GE Healthcare, Little Chalfont, United Kingdom) equilibrated with 20 mM Tris-HCl pH 8.0, 1 mM EDTA buffer purged with argon. THP was added to drive the reduction to completion as DTT stocks will contain trace amount of DTT_OX_. Protein concentrations were determined by absorbance at 280 nm with Nanodrop ND-1000 spectrophotometer (Thermo Fisher Scientific (Waltham, Massachusetts, USA)). To prepare a mixture for oxidizing conditions both homodimers (0.43 mg/ml of iOPRT^C44^ and 0.37 mg/ml OPRT^C92^) were incubated with 100 μM reduced DTT (the concentration of reduced DTT was determined with Ellman’s reagent, 5,5’-dithiobis-(2-nitrobenzoic acid)) in 20 mM Tris-HCl buffer pH 8.0, 1 mM EDTA for 20 minutes. Dimers were mixed and the oxidation was initiated by the addition of 10 mM oxidized DTT. Conditions for reducing conditions were prepared by incubating each homodimer separately with 2 mM of reduced DTT and 1 mM of THP for 20 minutes before mixing the homodimers together. Mixtures of both homodimers under reducing and oxidizing conditions were incubated at 30 °C and aliquots were taken at different time point to measure the activity. A 15% excess of iOPRT^C44^ was used over OPRT^C92^ since the disulfide bond formation was observed not to be complete and some residual activity would remain even with prolonged incubation in oxidizing conditions.

Activity measurement was based on the decrease of orotate absorbance at 290 nm as it is converted to orotidine monophosphate by the OPRT enzyme. The assay mixture was composed of 250 μM phosphoribosyl pyrophosphate (PRPP), 200 μM orotate and 5 mM MgCl_2_ in 20 mM Tris-HCl pH 8.0 buffer. An aliquot of dimerization reaction mixture (under oxidizing or reducing conditions) was taken at different time points, diluted ∼200 fold and 5 μl of diluted mixture was added to 145 μl of assay buffer, resulting in ∼6500-fold dilution overall. Absorbance at 290 nm was measured using a Perkin-Elmer Lambda 35S (Perkin-Elmer Inc., Waltham, Massachusetts) spectrophotometer in a temperature-controlled cuvette (30 °C). Initial rates were measured as decrease in absorbance over time and were obtained by linear or non-linear regression. Disulfide bond formation was assayed by quenching the samples with excess N-ethylmaleimide relative to DTT thiol groups and analyzing dimer/monomer ratios of the samples by non-reducing SDS-PAGE.

### Oxidation by *E. coli* Trx1

Reduced iOPRT^C44^ and OPRT^C92^ were prepared by incubating concentrated protein solutions in 50% glycerol for 30 minutes with THP/DTT_RED_ and desalting on a NAP5 column equilibrated with 20 mM Tris-HCl pH 8.0, 1 mM EDTA saturated with argon in argon atmosphere. 16.5 μM of each homodimer (33 μM of both homodimers) were mixed with 50 μM of *E. coli* thioredoxin 1 (Sigma-Aldrich GmbH, St. Louis, Missouri, USA), oxidized DTT or just adding Tris-HCl buffer as a control. Reaction was allowed to occur for 3 h at 30 °C and aliquots were taken at different time points. Each aliquot was added to 10% trichloroacetic acid, which decreases the pH value of the solution and precipitates proteins, thus the reaction should be inhibited. Precipitated proteins were pelleted by centrifugation at 13000 g at 4 °C for 1 h. Supernatant was removed, and the pellet was resuspended in 125 mM Tris-HCl, 4% SDS, 20% glycerol buffer containing bromocresol purple and 10 mM N-ethylmaleimide. 1 M Tris-HCl pH 8.0 was added to the sample until the color of bromocresol blue shifted from yellow to dark blue, indicating that the pH of the solution became optimal for N-ethylmaleimide to modify the free thiol groups. Samples were incubated for at least 15 minutes, heated to 95 °C and analyzed on acrylamide gels. Band intensities of oxidized and reduced forms were quantified with ImageJ software and plotted using the R programming language.

### Redox potential determination

Redox potentials were determined for iOPRT^C44^/ OPRT^C92^, OPRT^C44*^/OPRT^C92*^ and OPRT^C44^/^C92*^ disulfide bonds. The proteins were prepared for the reaction in the same way as for the thioredoxin experiment, although argon atmosphere was omitted, since it was observed that 1 mM EDTA and argon purged buffers are sufficient to prevent disulfide bond oxidation by atmospheric oxygen. Furthermore, 20 mM Tris-HCl pH 8.0 buffer was substituted with 50 mM sodium phosphate, pH 7.25. A range of redox buffers was prepared, with oxidized:reduced DTT ratios (mM): 50:0.5, 25:0.5, 12.5:0.5, 5:0.5, 2.5:0.5, 1:0.5, 0.5:0.5, 0.5:1, 0.5:2.5, 0.5:5, 0.5:12.5. Each homodimer was mixed at 0.4 mg/ml with the same amount of its disulfide bond partner and 0.8 mg/ml of double mutant (OPRT^C44/C92*^) and were incubated for 3 h at 30 °C. The reaction was quenched the same way as for the thioredoxin experiment and the samples were analyzed on acrylamide gels and quantified with ImageJ software.

### Differential scanning calorimetry

Prior to determination of the thermal profiles, homodimers were reduced with THP/DTT_RED_ mixture as described above and desalted. The protein concentration for differential scanning calorimetry experiments was 0.6 mg/ml (∼25 μM). Full oxidation of the disulfide bond in heterodimer mixtures (0.6 mg/ml of total protein) would require large amounts of oxidized DTT (50 mM), which might not be removed with NAP5 column and interfere with the measurements. This was indeed observed. DTT_OX_ was instead removed by micro-dialysis of a heterodimer mixture after 3 h of incubation at 30 °C. About 150 μl of protein sample was subjected to 3 rounds of dialysis in 250-350-fold excess of phosphate buffer at room temperature. Full oxidation was confirmed by SDS-PAGE.

### pKa determination

pKa values for OPRT^C44^ and OPRT^C92*^ variants were determined by analyzing their thiolate reactivities with iodoacetamide (IAM) at different pH values. 0.4 mg/ml (16.5 μM) of each variant reduced as previously. Proteins (desalted in 20 mM Tris-HCl buffer) were incubated in buffers with relatively high ionic strength to prevent electrostatic influences on thiolate reactivities and different pH values containing 1 mM of IAM for 1.5 minutes and the reaction was stopped by addition of trichloroacetic acid to 10%. Precipitated samples were incubated at 4 °C for at least 15 minutes, centrifuged at 16000 g and resuspended in 125 mM Tris-HCl, 4% SDS, 20% glycerol, bromocresol purple and 2 mM mPEG-maleimide buffer. pH was increased to ∼8 by addition of 1 M Tris-HCl pH 8.0 buffer and the modification of free thiol groups with mPEG was allowed to continue for 30 minutes. The following buffers containing 100 mM NaCl, 1 mM EDTA and 100 mM of buffering compound were used to make a pH scale: 5 (sodium acetate), 6 (MES), 7 (sodium phosphate), 8 (Tris-HCl), 8.5 (sodium tetraborate), 9 (sodium carbonate), 10 (CAPS), 11.3 (sodium phosphate). Samples were analyzed on 15% SDS-PAGE, two independent experiments were performed for each pH value for each of the variants. The k_obs_ values were obtained by least square regression of exponential decay of unmodified thiol and k_S-_ and pK_a_ values were obtained by non-linear least square approximation of the experimental data to the equation:

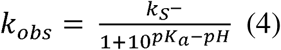

### Methods relating to yeast experiments

The *pyrE* cDNA was codon-optimized for yeast expression, and DNA encoding the two variants, OPRT* and OPRT^C44/C92*^, along with a C-terminal myc-tag, was synthesized and cloned into pDONR221 by Genscript. Using Gateway cloning (Invitrogen), these variants were then transferred into the destination vector pAG415GPD-ccdB (Addgene plasmid # 14146) (Alberti et al., 2007) for yeast expression. The BY4741 strain was used as the wild-type strain. All yeast knockout strains were obtained from the Euroscarf collection. Before transformation, yeast cells were cultured in Yeast Extract-Peptone Dextrose (YPD) medium (2% D-glucose, 2% tryptone, 1% yeast extract). After transformation with *pyrE*-encoding plasmids, cells were cultured in synthetic complete (SC) medium lacking leucine (2% D-glucose, 0.67% yeast nitrogen base without amino acids, 0.2% dropout supplement (US Biological), (2% agar)). Expression plasmids were transformed into yeast using the method described (Gietz and Schiestl, 2007).

For cultures used for protein extraction, the relevant yeast strain was inoculated into 20 ml SC-LEU medium and cultured overnight until reaching mid-exponential phase. For pertubation of the intracellular redox environment, exponentially growing yeast cells were incubated for 1 hour with varying concentrations of DTT or DPS. Specifically, DTT was dissolved in DMSO at 250 mM and was added to cultures at final concentrations of 0, 0.5, 1 or 10 mM. DPS was dissolved in DMSO at 20 mM and was added to cultures at final concentrations of 0, 0.05, 0.5 or 5 µM.

For yeast protein extraction, 100-125 × 10^6^ cells were harvested from exponential phase cultures by centrifugation (1,200 g, 4 °C, 5 min). All subsequent steps were performed on ice. The cell pellet was washed in 25 ml water (1,200 g, 4 °C, 5 min), and resuspended in 1 ml 20% TCA. After centrifugation (4,000 g, 4 °C, 5 min), the supernant was removed and the pellet was resuspended in 200 µl 20% TCA. This suspension was transferred into a screw-cap tube containing 0.5 ml acid-washed glass beads (Sigma-Aldrich). Cells were lysed using a Mini Bead Beater (BioSpec Products) with three cyles of 10 seconds each, interspersed with 5-minute incubations on ice. Next, 400 µl 5% TCA was added, and the tubes were punctured at the bottom with a needle. The tubes were placed on top of a 1.5 ml Eppendorf tube inside a 15 ml Falcon tube. Lysates were then transferred to the Eppendorf tubes by centrifugation (1,000 g, 4 °C, 5 min). The collected lysate was then centrifuged again (10,000 g, 4 °C, 5 min), and the resulting pellet was washed twice in 0.5 ml 80% acetone (10,000 g, 4 °C, 5 min) before being air-dried. The dried pellet was resuspended at room temperature in 200 µL 1.5x SDS sample buffer (prepared by diluting 4x SDS sample buffer: 250 mM Tris/HCl, 8% SDS, 50% glycerol, 0.05% bromophenol blue, pH 6.8) containing 10 mM NEM. If necessary, the pH was adjusted with 1 M Tris (pH 8.0) until the solution turned dark blue, then incubated at room temperature for 15 minutes. Finally, each sample was split in two; one aliquot was reduced using 3 µl 1 M DTT (to a final concentration of 29 mM DTT), and all samples were incubated at 90 °C for 5 minutes and were then stored at -20 °C prior to SDS-PAGE analysis.

Protein extracts were resolved by SDS-PAGE on 12.5% acrylamide gels. Following electrophoresis, proteins were transferred to 0.2 µm nitrocellulose (Advantec) membranes for Western blotting. Membranes were blocked in PBS containing 5% fat-free milk powder, 5 mM NaN_3_ and 0.1% Tween-20. Blots were then probed with an anti-myc primary antibody (Chromotek, 9e1) and an HRP-conjugated anti-rat secondary antibody (Invitrogen, 31470). Band intensities were quantified using Image Lab software (Bio-Rad).

## Results

### Design of a disulfide bond

*E. coli* orotate phosphoribosyl transferase (OPRT) is a homodimer composed of 23.6 kDa subunits related by 2-fold symmetry. We used geometry-based algorithms (MODIP (Dani et al., 2003), Disulfide by Design (Dombkowski, 2003), BridgeD (Pellequer and Chen, 2006) on an X-ray structure of *E. coli* OPRT (PDB ID: 1ORO) (Henriksen et al., 1996) to locate putative sites for the insertion of an inter-molecular disulfide bond. All of these algorithms predicted that the cysteine substitution of opposing R44 and D92 residues would result in a left-handed disulfide bond of the most favorable geometry (Figure 1). In general, it is worth noting that in a homodimer interface the opposing residues will, except for rare cases, be placed at different positions in the sequence.

**Figure 1.**
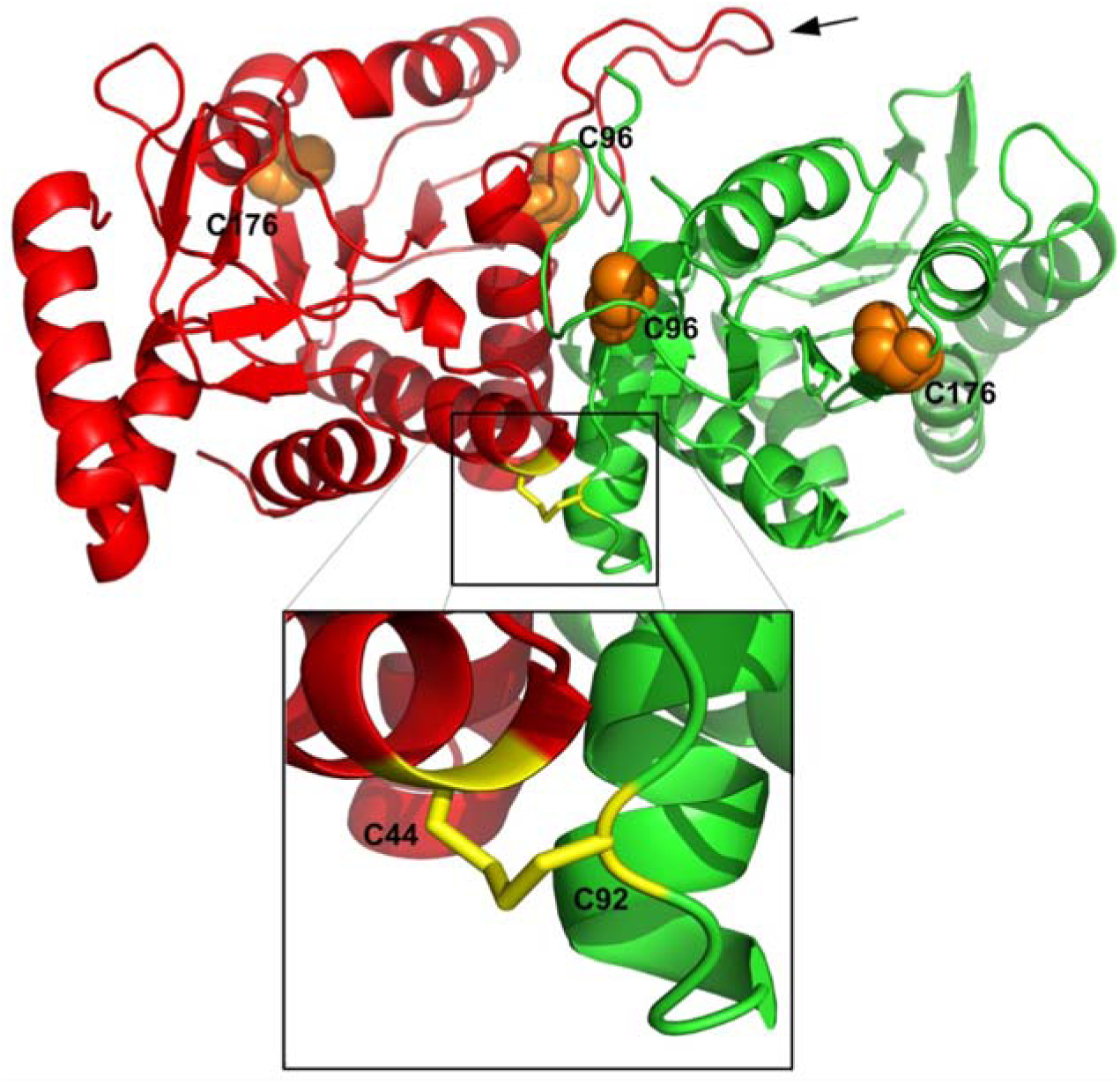
OPRT with native cysteine residues shown as space filling spheres and the designed asymmetric disulfide shown as a stick model (zoomed in insert). The two subunits are shown in green and red, respectively. The active site is comprised of a binding pocket on one monomer covered by a loop (shown with arrow) on the other in a “yin-yang”-like configuration. The structure was rendered based on pdb:1ORO. The shared active site means that hetero-dimers of a wild-type subunit and a subunit lacking critical residues in both the binding pocket and the loop will be inactive.

Small perturbations of the protein structure introduced by applying CONCOORD (de Groot et al., 1997) resulted in the C44-C92 bond still being predicted by these algorithms, indicating that the disulfide cross-link would not result in severe reduction of conformational entropy. In addition, the predicted disulfide a) interconnects a helix with a coil, a feature often observed in naturally occurring disulfide bonds (De Simone et al., 2006), b) is not located in the immediate vicinity of an active site loop, the mobility of which is required for the catalytic activity of the protein, and c) is not in a region of the protein known to have any direct functional relevance (Henriksen et al., 1996).

To test whether we could obtain a stable disulfide bond, we generated a number of OPRT variants where residues Arg44 and Asp92 were substituted by cysteine (Table 1). Wild-type OPRT also contains two unpaired cysteine residues in each subunit (C96 and C176) which we feared might interfere. Thus, we also made R44C and D92C variants in which the endogenous cysteine residues, C96 and C176, were replaced by Ala, for simplicity here denoted OPRT^C44*^ and OPRT^C92*^. A variant with the potential to form two inter-subunit disulfides, but lacking endogenous cysteines, was also generated (OPRT ^C44/92*^). Finally, a catalytically inactive C44-variant iOPRT^C44^ containing mutations in both parts of the inter-subunit shared active site was also made (Table 1). Variant dimers differing in subunits by at least one residue substitution are here referred to as *heterodimers*.

**Table 1.** List of OPRTase variants with corresponding substitutions.

| **OPRTase variant** | **Characteristics** | **Substitution(s)** |
| --- | --- | --- |
| OPRT<sup>C44</sup> | Single Cys variant in active subunit | R44C |
| OPRT<sup>C92</sup> | Single Cys variant in active subunit | D92C |
| OPRT<sup>*</sup> | Variant lacking endogenous cysteine residues | C96A/C176A |
| OPRT<sup>C44*</sup> | Single Cys variant in active subunit lacking endogenous cysteine residues | R44C/C96A/C176A |
| OPRT<sup>C92*</sup> | Single Cys variant in active subunit lacking endogenous cysteine residues | D92C/C96A/C176A |
| OPRT<sup>C44/C92*</sup> | Double Cys variant in active subunit lacking endogenous cysteine residues | R44C/D92C/C96A/C176A |
| iOPRT<sup>C44</sup> | Single Cys variant in an inactivated subunit | R44C/D124N/D125N/R99G/<br>K100S/K103S/H105G |

### Disulfide bond formation

OPRT variants with D92C and R44C substitutions were produced as described in Materials and Methods by Ni^2+^-affinity chromatography yielding 40-60 mg/liter of cell culture of pure hexahistidine C-terminally tagged protein for each variant in a single step (Figure S1). Incubation of equal amounts of each variant in buffers with different GSH/GSSG ratios and quenching the reaction with excess of N-ethylmaleimide resulted in a nearly complete shift of a band from ≈24 kDa (monomer) to ≈50 kDa (dimer) on non-reducing SDS-PAGE (Figure 2) indicating the formation of an intermolecular disulfide bond. Even at large molar excess of GSH the majority of the protein remained in the dimer state. Thus, it appeared that the introduced disulfide bond would be impossible to analyze with the GSH redox couple (E°’ = -240 mV (Aslund et al., 1997)) as even the best commercial preparations of GSH contain a significant contamination of GSSG (Winther and Thorpe, 2014). DTT, on the other hand, has a standard biochemical redox potential (E°’) of about -330 mV. Furthermore, unlike GSH, mixed disulfides with DTT are unlikely to accumulate to a significant extent (Creighton, 1986), simplifying the determination of redox potentials.

**Figure 2.**
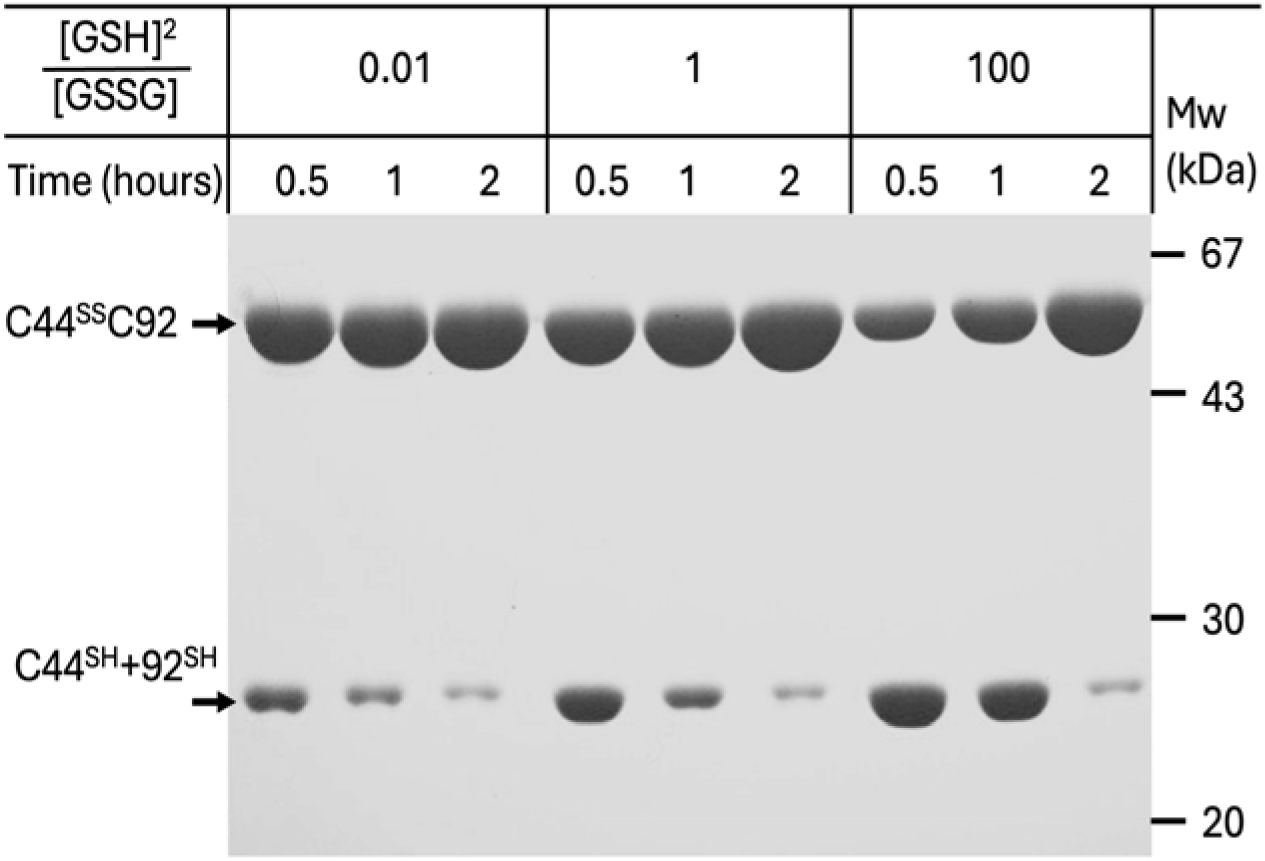
Formation of intermolecular disulfide bonds over time upon incubation of OPRT^C44^ and OPRT^C92^ with glutathione redox buffers. Disulfide bond formation is followed by non-reducing SDS-PAGE over time. Disulfide bond formation (conversion of C44^SH^+C92^SH^ to C44^SS^C92) is relatively rapid and is almost complete even under the most reducing conditions.

Using DTT, the reversibility of the disulfide bond formation could be demonstrated, as the ratio between oxidized and reduced OPRT was dependent only on the redox status of the buffers and independent of whether the starting point was oxidized or reduced OPRT (SI-Figure S2).

If the thiol groups were not quenched with N-ethylmaleimide prior to heating in 2% SDS sample buffer, the protein would migrate exclusively as a ≈24 kDa band. This can be explained by a thiol disulfide exchange reaction with either of the two Cys residues present in the wild-type protein (C96 or C176), resulting in isomerization from an inter- to an intramolecular disulfide bond. This interpretation was confirmed as introducing the C96A/C176A substitutions (to form OPRT^C44*^and OPRT ^C92*^) resulted in migration as a dimer even when the N-ethylmaleimide was omitted. Although homodimers of OPRT^C92^ and OPRT^C44^ were expected to be sterically unfavorable a small amount of C92-C92 and C44-C44 disulfide bonds was found if exposed to atmospheric oxygen for prolonged periods. Homodimers containing these disulfide bonds, however, exhibited different mobilities on an SDS-PAGE, and these disulfide bonds would be rapidly isomerized to more favorable C44/C92 upon the mixing of heterodimers.

### Redox potential determination

Despite the abundance of attempts to introduce artificial structural disulfide bonds into proteins, redox properties of the introduced disulfides are very rarely determined. As we wished to gain more insight in the stabilities of the introduced structural disulfide bond, we next determined the redox potential of the C44-C92 disulfide couple. Thus, mixtures of equal amounts of OPRT ^C44*^ and OPRT^C92*^ were equilibrated in phosphate buffer pH 7.25 with varying ratios of DTT_RED_/DTT_OX_ (25-0.01) present in large excess over protein. The extent of disulfide bond formation in each buffer was evaluated by non-reducing SDS-PAGE as described above.

Non-linear least square regression to equation 2 above was used to acquire K_OX_ from experimental data. A fraction of protein migrated as a 24 kDa band even under the most oxidizing conditions, indicating that the ratio between the two cysteine variants was not perfectly balanced. This fraction was subtracted before making K_OX_ calculations. The resulting K_OX_ values for the two variants with and without the native cysteines, iOPRT^C44^+OPRT^C92^ (K_ox_ = 0.286± 0.023) and OPRT^C44*^+OPRT^C92*^ (K_ox_ = 0.275±0.025), were identical within the experimental error (Figure 3C).

**Figure 3.**
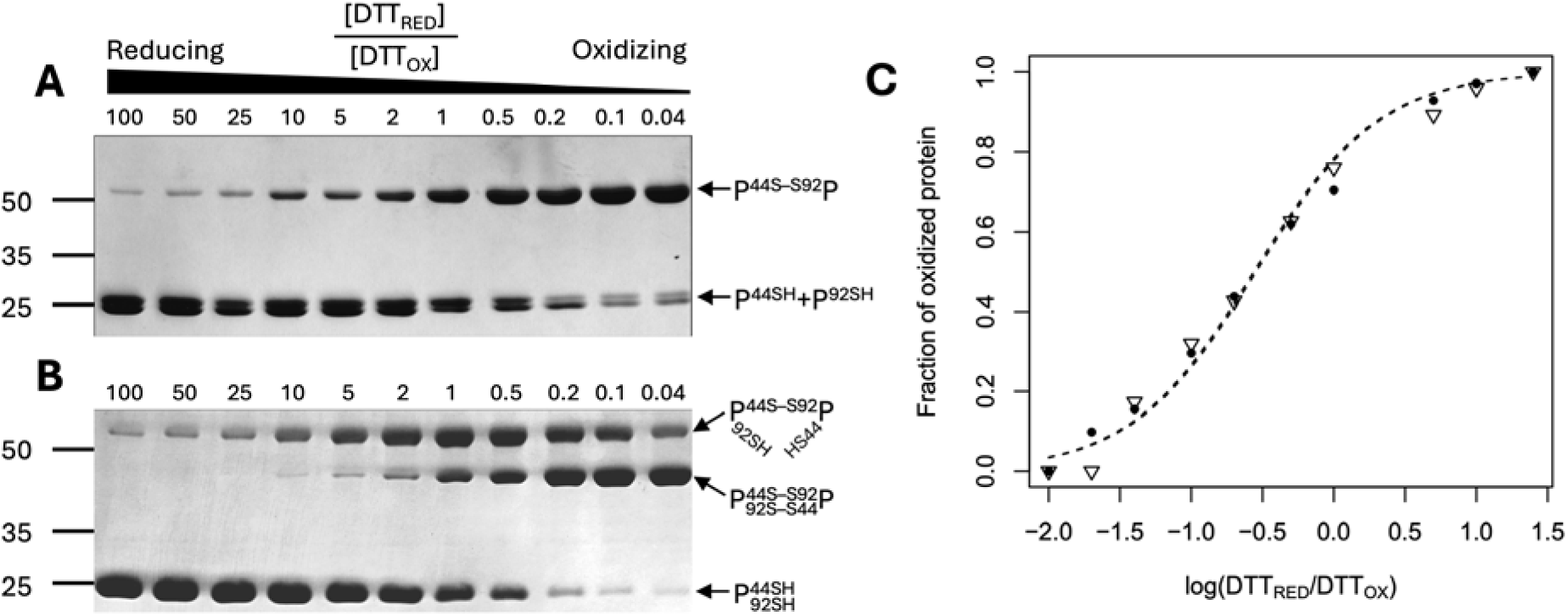
Redox titration of artificial disulfide bonds. (**A**) Non-reducing SDS-PAGE showing OPRT^C92*^+ OPRT^C44*^ after exposure to varying DTT_RED_/DTT_OX_ ratios. The disulfide-bonded oxidized heterodimer running at slightly above 50 kDa and the reduced forms at 25 kDa where the double band is likely due to the difference in charge between the two variants. (**B**) Same experiment showing OPRT^C44/C92*^ after exposure to varying DTT_RED_/DTT_OX_ ratios. From the top of the gel, the three forms represent single disulfide-bonded, double-disulfide-bonded and reduced forms, respectively. (**C**) Fraction of disulfide formation was determined from gel scans as in A for protein-variants with or without the native cysteine residues ((⍰) OPRT^C44^+ OPRT^C92^ and (●) OPRT^C44*^+ OPRT^C92*^, respectively). Data were fitted to a two-state redox model as described in Materials and Methods.

We also analyzed disulfide bond formation in the OPRT^C44/C92*^ homodimer. As expected, this variant exhibited the formation of a double-disulfide-bonded form, which can be seen in Figure 3B as the band with a migration rate between the reduced and single-disulfide bonded form on the gel. This band only gains prominence at more oxidizing conditions, in effect extending the dynamic range in redox sensing applications.

Adopting an E°’ of -330 mV for DTT the values of E°’ for disulfide bond formation are identical in OPRT^C92^+iOPRT^C44^ and OPRT^C92*^+OPRT^C44*^, at 0.314±0.001. It should be noted that the reactions were carried out at a pH value slightly above 7 and that conditions were therefore strictly speaking not standard. However, with a *pK_a_*well above 7.5 both for DTT and the involved protein thiols (see Figure 6), K_OX_ values should not vary considerably depending on pH.

### Disulfide bond formation kinetics

Mixtures containing equal amounts of OPRT^C92^ and OPRT^C44^ variants under oxidizing and reducing conditions exhibited specific enzyme activities very similar to that of wild-type OPRT (Table 2), indicating the absence of gross structural perturbations, which could have been caused by the side-chain substitutions or by the formation of the disulfide bond. The activity of the reduced form of the double mutant OPRT^C92/C44*^ was 7-fold lower than the wild-type enzyme, but upon oxidation of both disulfide bonds the enzyme activity was fully restored.

**Table 2.**
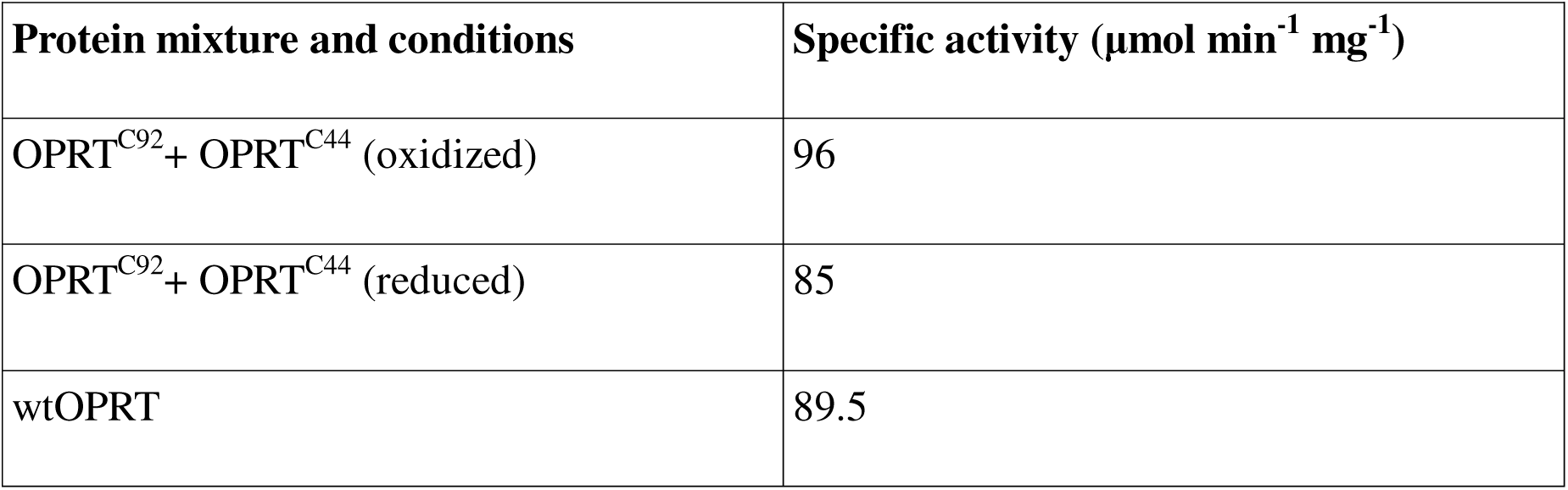
Specific activities of the variants under different redox conditions.

|  |  |
| --- | --- |
| Protein mixture and conditions | Specific activity ($\mu\text{mol min}^{-1} \text{mg}^{-1}$) |
| OPRT<sup>C92</sup>+ OPRT<sup>C44</sup> (oxidized) | 96 |
| OPRT<sup>C92</sup>+ OPRT<sup>C44</sup> (reduced) | 85 |
| wtOPRT | 89.5 |

To evaluate the kinetics of disulfide bond formation, an active site complementation experiment was performed similarly as done previously with OPRT from the *Salmonella typhimurium* enzyme which has 99% sequence identity to that from E. coli (Ozturk et al., 1995). The OPRT dimer contains two shared active sites, which are comprised of a substrate binding site in one subunit and a covering loop from the other. Closure of the flexible loop excludes solvent water from the active site and provides K103 which is essential for catalysis (Ozturk et al., 1995). An inactive OPRT^C44^ variant (iOPRT^C44^) was made by introducing active site substitutions in both the binding site and the flexible loop as described in Materials and Methods. Because of the yin-yang shared-active site structure of OPRT, a heterodimer between such an active-site mutant monomer and a wild-type monomer will be inactive (Ozturk et al., 1995). Thus, since only the OPRT^C92^ homodimer is active, random exchange of subunits of an active OPRT^C92^ dimer and iOPRT^C44^ variant would result in a time-dependent decrease of activity, reaching 50% of initial activity when the two different modified subunits are randomly mixed in the absence of disulfide bond formation. However, on shift to oxidizing conditions, the activity should over time approach zero, since the conversion to heterodimer would be driven by disulfide bond formation. This is indeed observed (Figure 4A). In the experiment shown in Figure 4A a 15% excess of the iOPRT^C44^ (inactive) over OPRT^C92^ (active) was used to compensate for incomplete disulfide bond formation under the experimental conditions applied (Figure 4B). A relatively high residual activity was seen even after prolonged incubation of heterodimer under oxidizing conditions.

**Figure 4.**
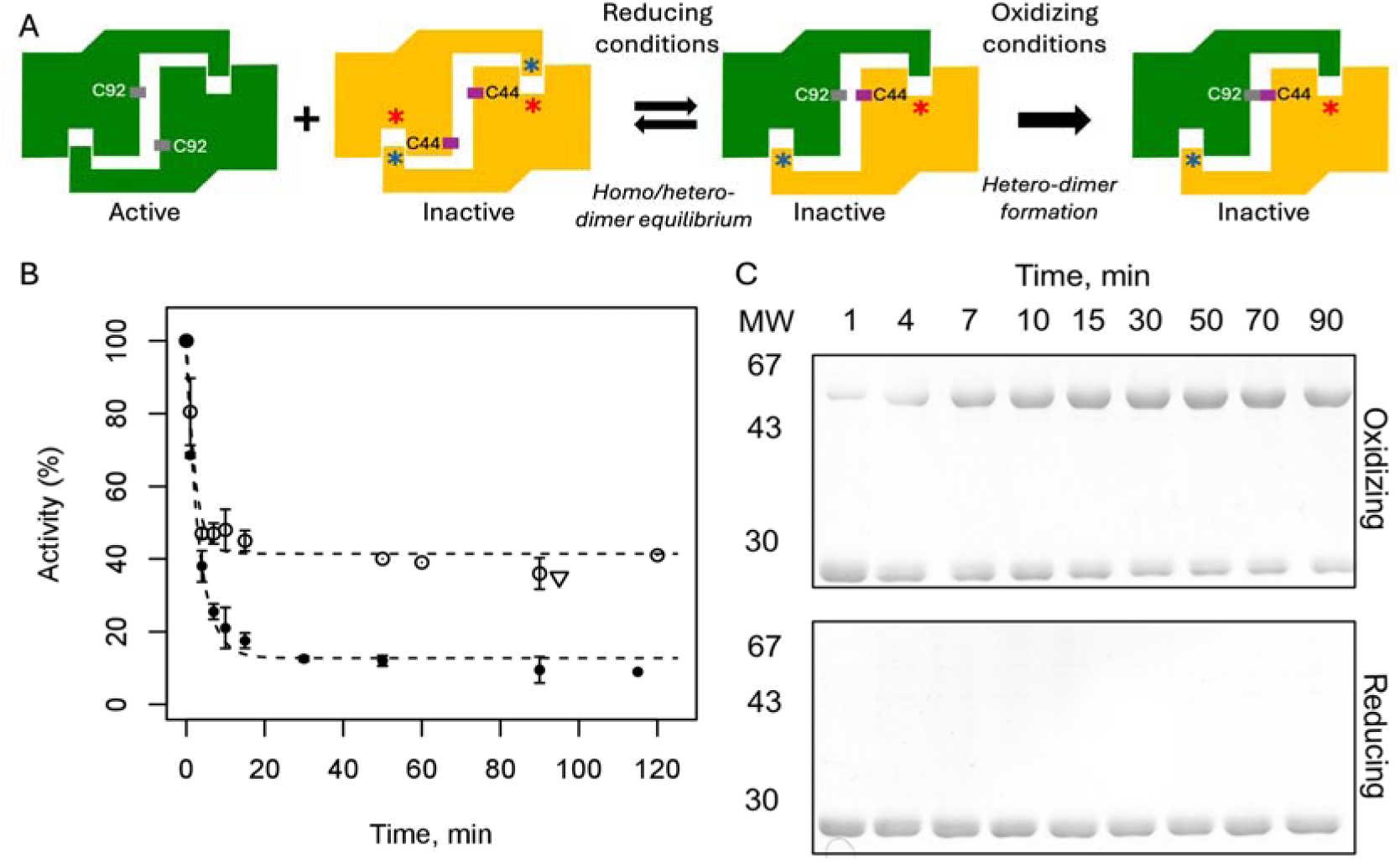
Kinetics of disulfide bond formation analyzed by active-site non-complementation. (A) OPRT is a homodimer with a shared active site comprising a substrate-binding groove and a covering loop. Heterodimers formed between an active (green) variant and a mutant (orange) variant containing disabling mutations in both the loop (blue star) and the groove (red star) are inactive. The active and inactive variants also contain C92 and C44 substitutions, respectively. Under reducing conditions, homo- and heterodimers distribute in a 1:1:2 ratio (green-green : orange-orange : green-orange). Thus, theoretically, mixing green and orange homodimers results in a 50% activity reduction under reducing conditions. However, under oxidizing conditions, disulfide bond formation drives the formation of heterodimers, resulting in a complete loss of activity. (B) Activity was measured over time after mixing of homodimers as described above. At the start of the reaction the OPRT^C92^ homo-dimer represents 100% activity. Negative active site complementation of iOPRT^C44^ and OPRT^C92^ under reducing (○) and oxidizing (●) conditions reduces activity to half by random association of active and inactive species due to formation of an oxidized heterodimer. The activity of this oxidized heterodimer is restored after 10-minute reduction with THP (⍰). Dotted lines indicate single exponential fit and error bars indicate one standard deviation. (C) Non-reducing SDS-PAGE as in figure 1. The formation of disulfide bonds in the OPRT^C92^ and iOPRT^C44^ heterodimer is followed over time under oxidizing (top gel) and reducing (bottom gel) conditions during the experiment. A small fraction of homodimer after 90 min is attributed to slight deviation from equimolar amounts of the two variants.

The rate-limiting step for the exchange of monomers in a dimer could, in principle, be dictated by either dissociation or association rates of the subunits. Ozturk et al. have shown that indeed the dissociation is limiting the exchange rate for OPRT monomers (Ozturk et al., 1995). Hence, if the disulfide bond formation is slower than the rate of dissociation, the rate of activity decay should be considerably different between oxidizing and reducing conditions since complete formation of the heterodimer is driven by disulfide bond formation. To compare rates, the activity decay is modeled as a first order reaction (Figure 4B, dashed lines). The rate constants for the decline in activity of reduced and oxidized mixtures do not differ substantially, being 0.45±0.21 min^-1^ for reduced disulfide and 0.3±0.085 min^-1^ for oxidized. Therefore, the rate of disulfide bond formation must be limited by dimer dissociation. These rate constants of disulfide bond formation are in good agreement with the ones calculated from the disulfide bond formation observed on acrylamide gels (0.22±0.09 min^-1^) (Figure 4C).

### Oxidation by thioredoxin

The observation that the redox potential for the C44/C92 disulfide was very low prompted us to test if the *E. coli* thioredoxin, *Trx1,* could oxidize this disulfide bond. Even though *Trx1* has one of the lowest redox potentials (-290 mV) determined for any CXXC active site redox catalysts (El Hajjaji et al., 2009), the oxidation of the artificial OPRT disulfide bond should still be thermodynamically favorable. To test this, 3-fold excess of oxidized thioredoxin was mixed with reduced iOPRT^C44^/OPRT^C92^ heterodimer under oxygen-free conditions at pH 8.0 and oxidation of the disulfide bond was followed over time by quenching with N-ethylmaleimide followed by non-reducing SDS-PAGE. DTT_OX_ was used as a low-molecular weight oxidant control. While no significant oxidation occurred in the absence of oxidized thioredoxin and DTT_OX_, addition of either of the components resulted in the oxidation to form a disulfide bond (Figure 5). No mixed disulfides were observed between OPRT monomers and Trx (Figure S3) showing, not surprisingly, that the half-life of this intermediate is short. In spite of possible steric hindrance, the rate of disulfide bond formation was significantly faster in the Trx-containing mixture compared to the DTT_OX_ mixture at identical concentrations of the oxidant. This demonstrates that oxidation of the disulfide bond is indeed catalyzed by Trx, albeit not at a high rate.

**Figure 5.**
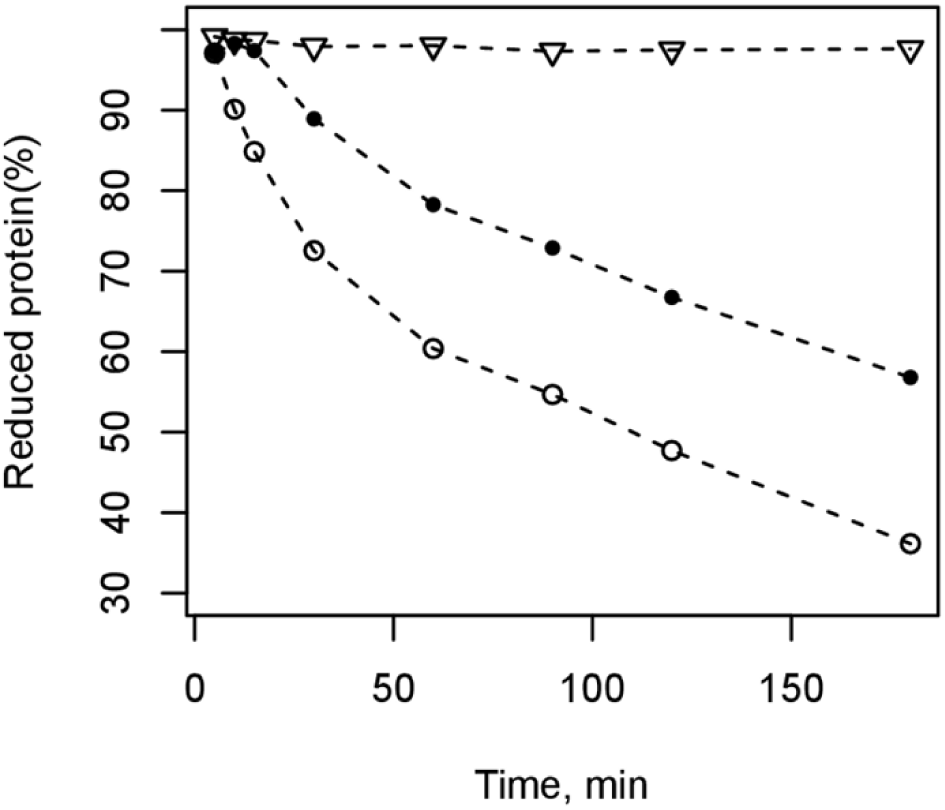
Kinetics of oxidation of OPRT^C92*^ and OPRTC^44*^ with thioredoxin. A 1:1 mixture of 16.5 μM of OPRT^C92*^ and OPRT^C44*^ was mixed with 50 μM of DTT_OX_ (●) or 50 μM *E. coli* thioredoxin 1 (○), respectively. A control experiment without addition of oxidant is also shown (⍰).

### pKa determination

In the OPRT^C44^ homodimer C44 is located at the N-terminal end of an α-helix, therefore the thiolate should be stabilized by a positive dipole of the helix but is, on the other hand, proximal to D92 on the opposing monomer. Inversely, C92 resides near a positively charged arginine on the opposing monomer.

The *pKa* values of the thiol groups in the OPRT^C44*^ and OPRT^C92*^ variants were determined using iodoacetamide as a thiol-modifying agent, essentially as described previously (Iversen et al, 2010). In this assay, the reactivity of iodoacetic acid is dependent on the protonation state of thiolate, and the *pK_a_* can thus be determined from the reaction rates at different pH values. The degree of iodoacetamide modification is determined by subsequent labeling with mPEG-maleimide, which increases the molecular weight of the unreacted protein by 2 kDa. The extent of modification can thus be visualized by analyzing samples by SDS-PAGE. While the thiol group of OPRT^C44*^ was fully modified with mPEG-maleimide (Figure S4), only ∼50% mPEG-maleimide modification was seen for the OPRT^C92*^ thiol even at low pH. Here it should be essentially nonreactive towards iodoacetamide and thus undergo full mPEG modification. Although we cannot explain this loss of reactivity, it can be reasoned that the same amount of the thiol was not participating in the reaction in all of the samples. Hence, the fraction of unreacted protein was subtracted, and the extent of modification was normalized.

The pK_a_ values obtained for C44 and C92 thiols were 9.14±0.05 and 8.12±0.1 respectively (Figure 6). This is consistent with the presence of R44 close to C92 in the opposite monomer, stabilizing the thiolate and lowering the pK_a_. On the other hand, the vicinity of the D92 has the opposite effect on C44. Maximum reaction rates, ks-values, on the other hand, were essentially the same at 1.5 M^-1^min^-1^ for both thiols.

**Figure 6.**
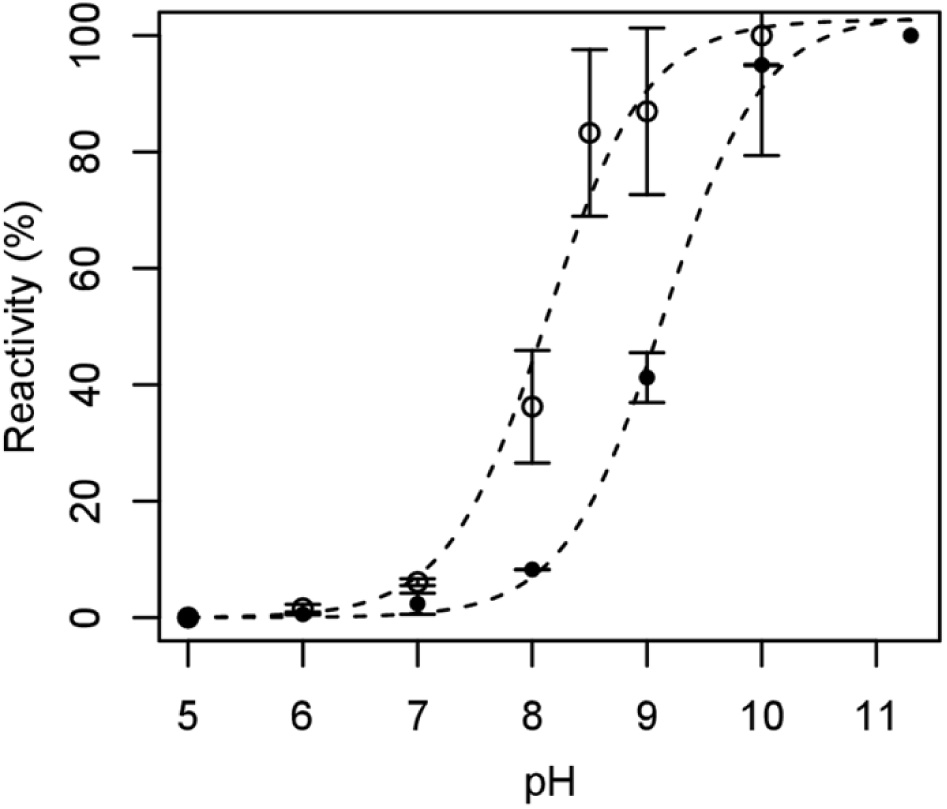
Determination of thiol pKa values for the two cysteines introduced. The pH dependence of the relative reaction rates of (○) - OPRT^C92*^ and (●) - OPRT^C44*^ thiols is shown. Dashed lines indicate the non-linear least squares estimation as described in the text and show a pKa of 8.1 for Cys92 and 9.1 for Cys44. Error bars indicate one standard deviation.

### Determination of global thermostability

The influence of the disulfide bond(s) on the protein stability was analyzed with differential scanning calorimetry (Figure 7; Table 3). One peak was observed for all the protein variants in a range of 30 to 90 °C (Figure 7), followed by a very intense heat emission which is indicative of protein aggregation. A twofold decrease or increase in protein concentration did not affect the position of the peak, suggesting that the dimer-monomer equilibrium is not part of the unfolding pathway. Instead, it is more likely that dissociation and unfolding occur as concerted reactions.

**Figure 7.**
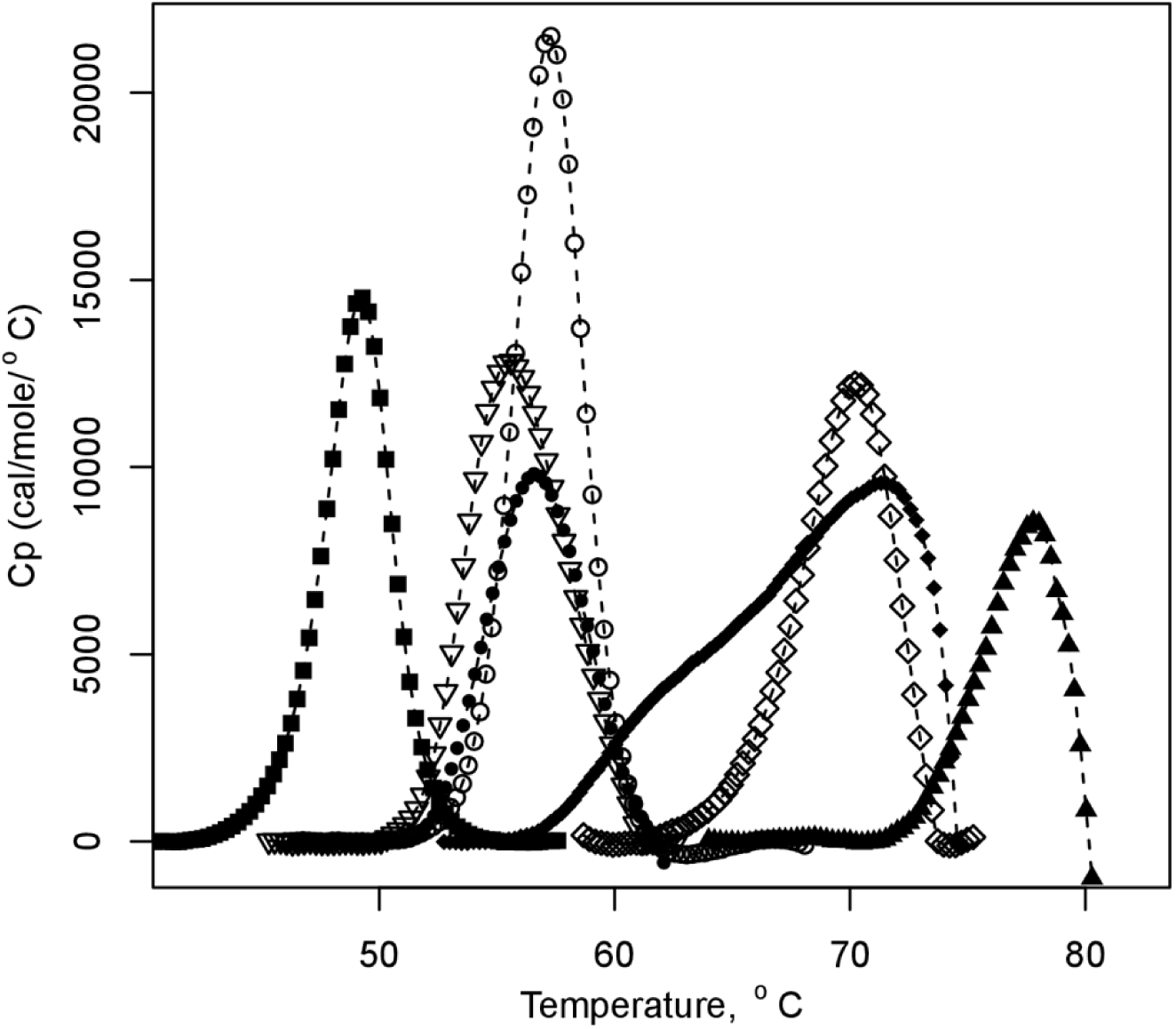
Thermal unfolding profiles of protein variants, iOPRT^C44^ (▪), OPRT^C44*^ (⍰), OPRR^C92^ (o), OPRT^C92*^ (●), wtOPRT (♦), OPRT^C92*^ + OPRT^C44*^ (oxidized, (◊)) and OPRT^C44/C92*^ (oxidized, (▴)). It can be seen, that oxidized OPRT forms are at least as stable as wild-type protein, however, reduced forms have decreased stability relative to wild-type.

**Table 3.** T_m_ values obtained from differential scanning calorimetry.

| Table 3. T<sub>m</sub> values obtained from differential scanning calorimetry |  |
| --- | --- |
| Variant | T<sub>m</sub> °C |
| OPRT<sup>C92</sup> | 57.1 |
| OPRT<sup>C92</sup> + iOPRT<sup>C44</sup> (oxidized) | 69.9 |
| OPRT<sup>C92*</sup> | 56.7 |
| OPRT<sup>C44*</sup> | 55.1 |
| OPRT<sup>C92*</sup> + OPRT<sup>C44*</sup> (oxidized) | 77.4 |
| OPRT<sup>C44/C92*</sup> (reduced) | 52.2 |
| OPRT<sup>C44/C92*</sup> (oxidized) | 78.6 |
| wtOPRT | 71.5 |

The heat denaturation of all variants, including wtOPRT, was irreversible, allowing us to obtain only T_m_ of denaturation. The wtOPRT melting profile displayed a shoulder (Figure 7), which could indicate dimer dissociation prior to unfolding. A mixture of OPRT^C92^ and OPRT^C44^ variants under reducing conditions displayed almost identical unfolding profiles as corresponding variants analyzed individually, thus only the stabilities of homodimers are presented in Figure 7. Notably, all the cysteine variants were less stable than the wild type-protein in the absence of a disulfide bond, but the thermal stability was restored to wtOPRT level, or above, upon formation of the disulfide bond. Interestingly, OPRT^C92/C44*^ with both disulfide bonds oxidized did not increase the stability substantially compared to oxidized OPRT^C92*^+OPRT^C44*^ having only a single disulfide. On the other hand, OPRT^C92/C44*^ was stabilized to the largest extent (26.4 °C) when compared to its reduced form.

### Disulfide bond formation in the yeast cytosol

To test whether the engineered disulfide bonds would also form *in vivo*, we codon-optimized the *pyrE* sequence for yeast expression and expressed two variants: OPRT* (with both endogenous cysteines mutated to alanine) and OPRT^C44/C92*^ in yeast. To assess disulfide bond formation, we extracted total protein using TCA followed by homogenization with glass beads. Before boiling in SDS, samples were treated with NEM to prevent disulfide bond exchange. As expected, OPRT* migrated as a monomer at ∼24 kDa (Figure 8A). In contrast, OPRT^C44/C92*^ migrated as three distinct bands, corresponding to the single-disulfide bonded dimer, the double-disulfide bonded dimer, and the monomer (Figure 8A). Quantifications of diluted OPRT^C44/C92*^ samples from yeast revealed that approximately 44% exist in the single-disulfide bonded form, 51% in the double-disulfide bonded form, and only 5 % in the monomer form (Figure S5). Based on the previous redox titrations (Figure 3B) and an E°’ of -330 mV for DTT, this corresponds to a cytosolic redox potential in wild-type yeast of approximately -300 mV.

**Figure 8.**
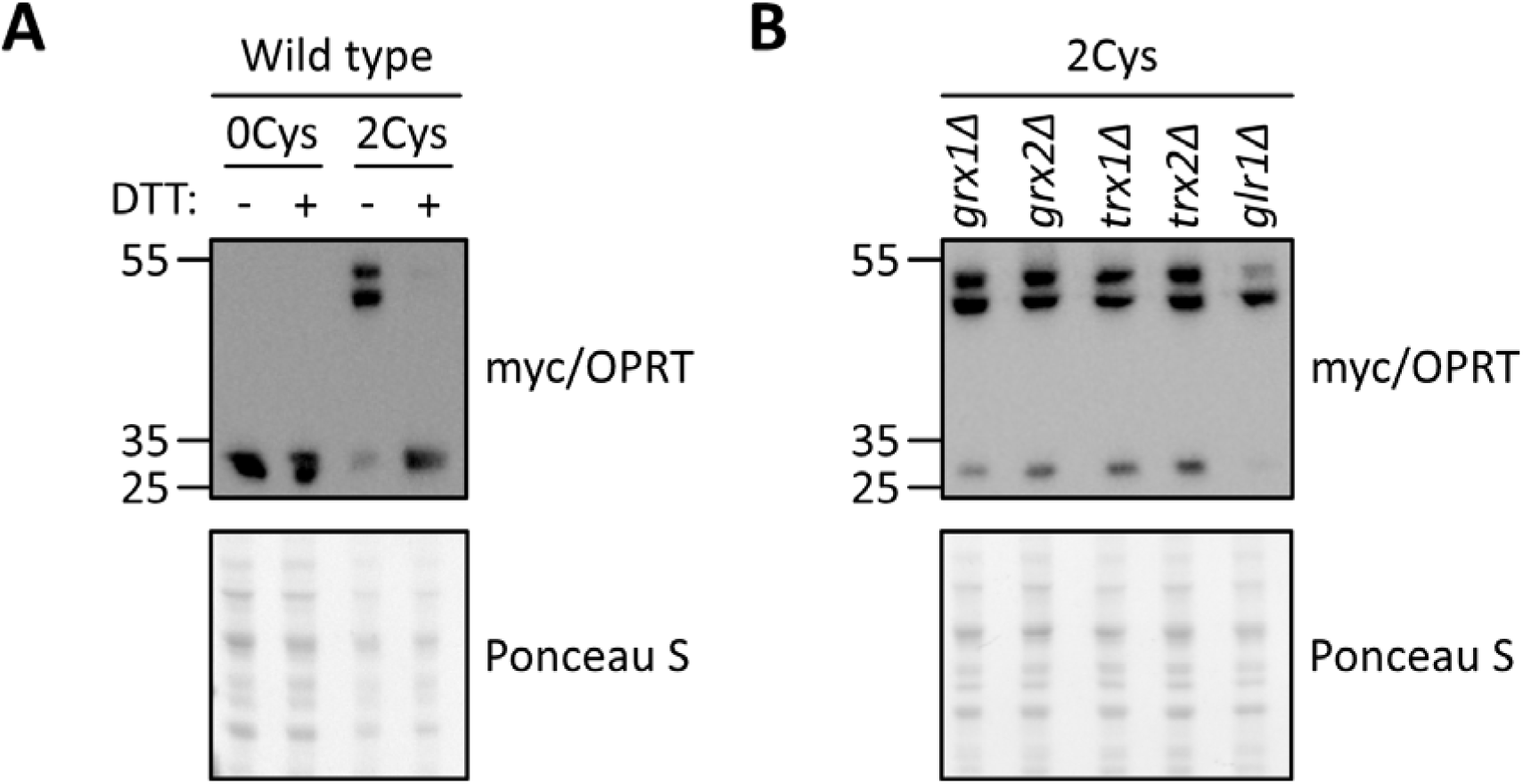
Formation of engineered OPRT disulfide bonds in the yeast cytosol. (A) Western blot showing that OPRT exists as a monomer in wild-type yeast expressing OPRT* (0Cys), whereas in yeast expressing OPRT^C44/C92*^ (2Cys), OPRT is present in three forms: monomeric, single-disulfide bonded, and double-disulfide bonded. (B) Western blot showing the distribution of OPRT across these three forms in the indicated redox knockout strains.

To further examine the disulfide bond *in vivo*, we treated yeast cells with the reducing agent DTT or the oxidizing agent 2,2-dipyridyldisulfide (DPS) before protein extraction. As expected, DTT treatment shifted the equilibrium of OPRT towards the reduced monomer form (Figure S6). In contrast, DPS treatment shifted the equilibrium towards the fully oxidized double-disulfide bonded dimer form (Figure S6). Collectively, these results show that the disulfides in the cytosolic OPRT dimer can be modulated using redox reagents.

Finally, we examined how yeast cellular redox systems influence the engineered cytosolic disulfide bond. We compared the proportions of OPRT in the three different forms (monomer, one disulfide, two disulfides) in a wild-type strain and five redox knockout strains: *grx1*Δ*, grx2*Δ*, trx1*Δ*, trx2*Δ *and glr1*Δ. Knockout of either of the glutaredoxins or thioredoxins only had a minor effect (Figure 8B), suggesting that the cell’s ability to reduce disulfide bonds is only slightly impaired. In the *glr1*Δ strain, where the cellular pool of reduced glutathione is lower (Ostergaard et al., 2001), the redox balance is shifted dramatically towards a more oxidized state. As a result, less OPRT is found in the fully reduced and single-disulfide forms, while the proportion of OPRT in the double-disulfide bonded form increases (Figure 8B). Collectively, these results suggest that the disulfide state is determined mainly by the glutatione/glutaredoxin system, supported by Glr1 activity, even when the disulfide is a substrate for thioredoxin.

## Discussion

We have successfully designed a very stable disulfide bond. The designed disulfide bond is expected to have dihedral angle values (**X**^1^: -64°/-70°, **X**^2^: 141°/-56.5°, **X**^SS^: -74°), which are well populated among naturally occurring structural disulfides (Dani et al., 2003; Petersen et al., 1999). Its conformation is predicted to have low dihedral strain energy (Marques et al., 2010), which was observed experimentally by a relatively low redox potential close to that of DTT. Although it is an intermolecular cross-link in a strict sense, a strong protein dimerization, as seen for OPRT, increases the effective concentration of reacting thiols by positioning them in a close spatial proximity. As a result, the low redox potential observed for the disulfide bond is more akin to intra-chain disulfides. The disulfide bond isomerized from intermolecular to intramolecular when the protein was heated in the presence of SDS. This behavior is expected, since the disruption of the dimer interface highly reduces the probability of encounter between C92 and C44 thiol groups. This diminishes the rate of the formation of this disulfide bond, while disrupting structure and increasing protein backbone mobility make it more likely that native C96 and/or C176 thiols engage in a disulfide bond.

As indicated by differential scanning calorimetry, the disulfide bond stabilized the OPRT^C44*^-OPRT^C92*^ dimer relative to the wild-type protein by approximately 6° C. However, geometry-based algorithms did not predict a large destabilization of the reduced form. Taking a thermodynamic cycle into consideration (Kim and Lin, 1989), the stability of the disulfide bond will be increased by decreasing the ΔΔG for unfolding of the reduced form of the Cys-containing species, as long as the native reduced state is more stable than the unfolded state. Therefore, it seems likely that geometry-based algorithms optimizing the oxidized state of the disulfide bond, but disregarding stabilities of half-cysteines, are likely to improve the stability of disulfide bonds themselves, but not the proteins into which they are introduced. The program for disulfide bond prediction, BridgeD (Pellequer and Chen, 2006), attempts to evaluate the destabilizing effect of substitutions. Nevertheless, the contribution of exposed salt bridges (as the putative one between R44 and D92) to the stabilization of proteins is controversial (Dong and Zhou, 2002) and the program BridgeD highly overestimated the free energy cost of C44/C92 substitutions, giving a disulfide bond relatively low ranking.

Substitutions had a profound influence on the stability of the reduced protein, destabilizing mutant variants by ∼16 °C for one introduced substitution and ∼20 °C when both substitutions are introduced. Although a steric “clash” is often proposed to explain the destabilizing effect of unsuccessfully introduced disulfide bonds (Mitchinson and Wells, 1989), it would in this case be unlikely, since a cysteine residue has only a slightly bigger volume (V_r_≈103 Å^3^) than Asp (V_r_≈91Å^3^) and is much smaller than Arg (V_r_≈150 Å^3^) (Creighton, 1994). Therefore, disruption of a salt bridge between Arg-Asp residues seems to be a more likely explanation for the reduced stability of the reduced form. These results demonstrate the potential complications associated with the introduction of disulfide bonds in proteins with the purpose of enhancing stability.

The intrinsic reactivities of C44 and C92 towards IAM were very similar (k_obs_ values 1.5 s^-1^), indicating, that the residues should be equally surface accessible. Lowered pK_a_ value for C92 (8.12 compared to about 9 for cysteines in peptides) could be explained by its proximity to the positively charged Arg residue in the homodimer of OPRT^C92*^ (Hansen et al., 2005). The pK_a_ value of C44 (9.14) is expected, since the N-terminus of an α-helix dipole could compensate for the destabilizing effect by the negatively charged D92 (Wada, 1976).

The surface accessibility of our engineered disulfide bond contrasts to naturally occurring disulfides, which are generally buried (Dani et al., 2003). The latter property hampers analysis of redox potential, and explains why most, if not all, well characterized disulfide bonds are from designs. Thus, the engineered C60-C173 disulfide of human carbonic anhydrase II has a redox potential of -337 mV (calculated from K_ox_ with DTT), although the oxidized protein is less stable than wild-type (Burton et al., 2000). Mitchinson and Wells introduced a series of disulfide bonds to subtilisin BPN’, several of which were extremely stable (Mitchinson and Wells, 1989).

To the best of our knowledge, the redox potential of OPRT^C44/C92*^ is the lowest determined for a designed *intermolecular* disulfide bond. As we show, the fact that it is intermolecular makes it an attractive tool for determination of intracellular redox status as the level of disulfide bond formation can readily be determined from immunoblots of non-reducing SDS-PAGE. An intramolecular disulfide bond engineered into Yellow Fluorescent Protein (rxYFP) had a redox potential of -265 mV as determined by equilibration with glutathione (Ostergaard et al., 2001). When expressed in yeast rxYFP was 10% oxidized at steady-state exponential cultures, while the equilibrium level was about 15% (as determined from pulse-chase labeling). This is because the newly synthesized rxYFP will only reach redox equilibrium over 5-10 min (Ostergaard et al. 2004). Thus, in exponentially growing cells a steady-state determination of redox status of a sensor will always slightly under-estimate the level of disulfide bond formation (Ostergaard et al. 2004). Given this, it appears that the estimate of thiol-disulfide redox conditions measured by OPRT^C44/C92*^ of about -300 mV is somewhat more reducing than the -289 mV determined in the yeast cytosol using rxYFP determined at equilibrium (Ostergaard et al. 2004). We note that while GFP-based probes are not substrates for thioredoxin, OPRT^C44/C92*^ is suggesting that thioredoxin may also play a role in reduction of OPRT^C44/C92*^.

## Abbreviations

DTT_OX_: oxidized dithiothreitol;
DTT_RED_: reduced dithiothreitol;
OPRT: orotate phosphoribosyltransferase;
THP: tris(hydroxymethyl)phosphine

## Acknowledgements

We thank Anne-Marie Lauridsen for expert assistance with the yeast assays.

## Funding

JRW was funded in part from the Novo Nordisk Foundation (https://novonordiskfonden.dk/) Grant nos. NNF19OC0058579 and NNF23OC0086187. R.H.-P. was funded by the Novo Nordisk Foundation (https://novonordiskfonden.dk/) (REPIN and NNF21OC0071057), the Danish Council for Independent Research (https://dff.dk) (10.46540/2032-00007B), and the LEO foundation (https://leo-foundation.org) (LF-OC-24-001531).

## Notes

### Competing Interest Statement

The authors have declared no competing interest.

### Summary of Updates

We have included additional data to show how the designed OPRT disulfides can be utilized to measure cytosolic redox conditions in yeast.

