## Supplementary information for "A highly stable engineered disulfide bond in the dimer interface of *E. coli* orotate phosphoribosyl transferase measures cytosolic redox conditions in yeast"

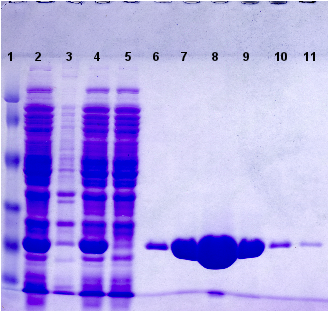

**Figure S1. Sample SDS-PAGE showing fractions from the purification of OPRT variants**

**from Ni-NTA column.** Lane 1 – Pierce Prestained Protein Molecular Weight Marker; lane 2 – sample taken after disruption of bacteria by sonication; lane 3 – pellet obtained after centrifugation of sonicated cells; lane 4 – soluble protein extract; lane 5 – column flowthrough after loading the protein extract; lanes 6-11 – 5 μL aliquots taken from 1-mL fractions of protein eluted with 100 mM imidazol buffer.

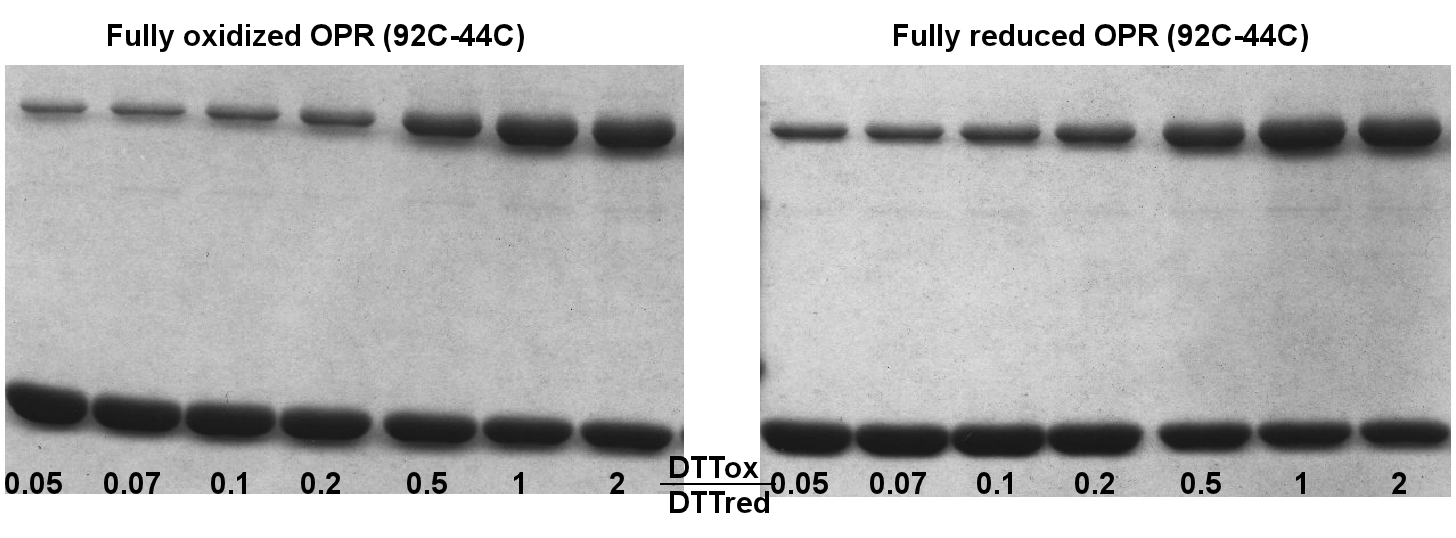

**Figure S2. Disulfide bond formation is reversible.** The distribution between monomer and dimer is the same whether starting with fully oxidized (left) or fully reduced (right) mixtures of OPRT^C44^ and OPRT^C92^ variants is reversible.

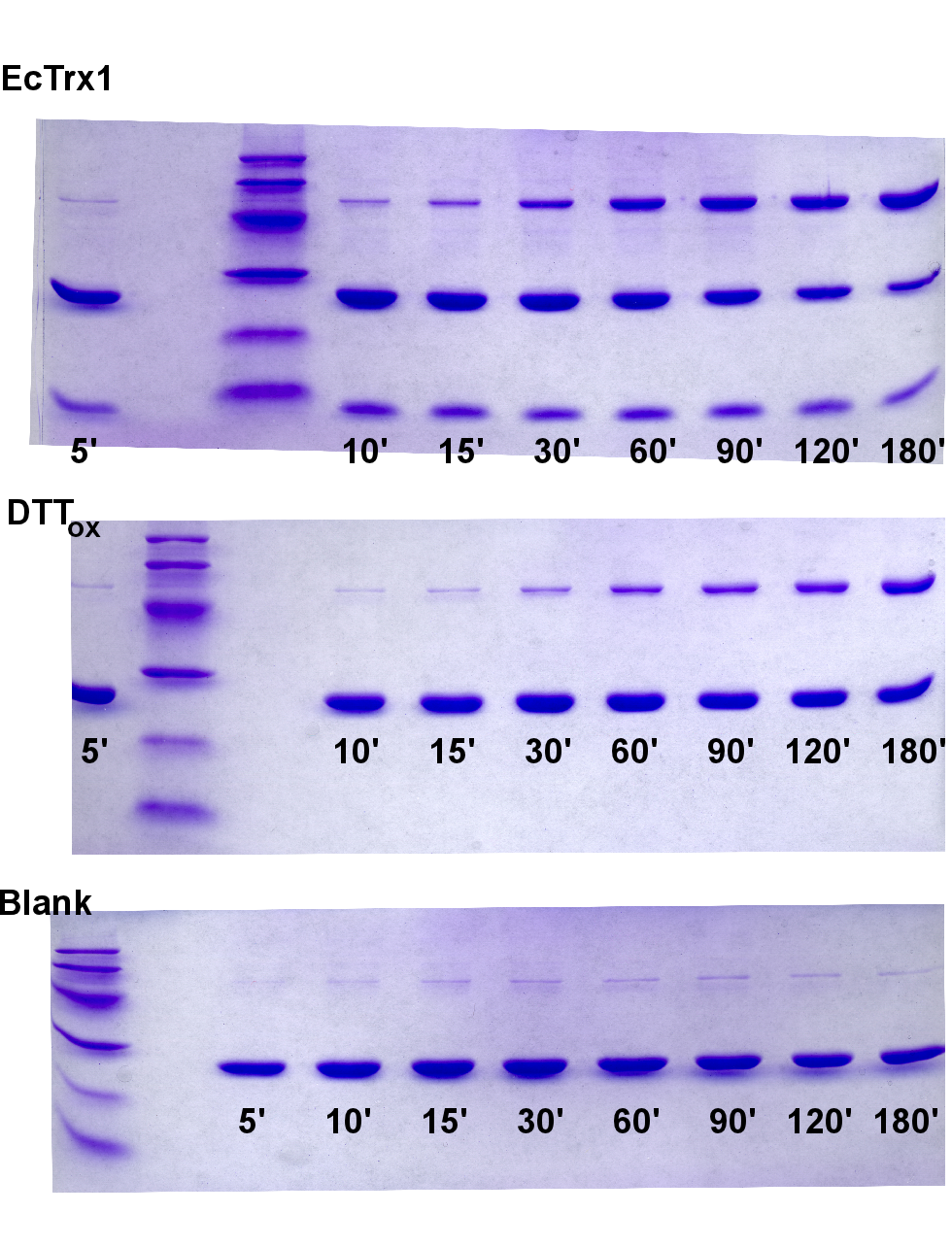

**Figure S3.** Raw non-reducing acrylamide gels used to quantify the rate of formation of the disulfide bond in iOPRT^C44^+OPRT^C92^ in the presence of thioredoxin (top), DTT_OX_ (middle) or buffer control (bottom). Numbers bellow the lanes indicate time in minutes from addition of oxidant. The fastest migrating band of the top gel is thioredoxin.

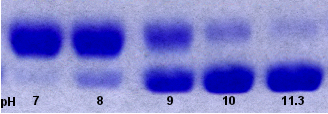

**Figure S4.** Representative SDS-PAGE showing reaction with IAM followed by PEGylation for OPRT^C44*^ at different pH values. Lane 1 – reaction at pH 7, lane 2 – pH 8, lane 3 – pH 9, lane 4 – pH 10, lane 5 – pH 11.3.

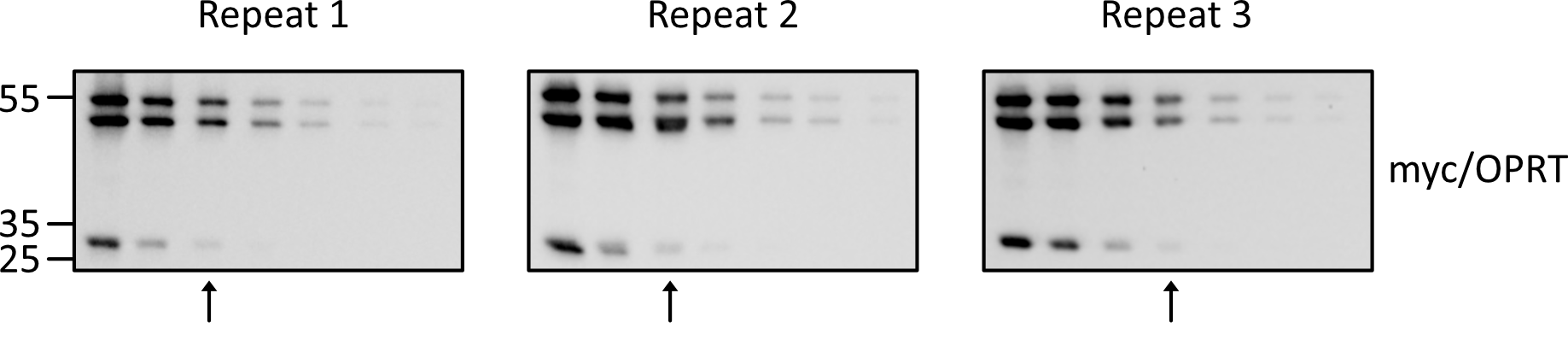

**Figure S5. Serial dilutions of OPRT^C44/C92*^ yeast extracts.** Western blots showing three repeats of three-fold serial dilutions of protein extracts from yeast cells expressing OPRT^C44/C92*^. Arrows indicate the lanes used for band intensity quantification, which determined the relative volumes of each band as a percentage of the total lane intensity. Based on quantifications from the three repeats, the mean proportions of the three bands are: 43.8% (±6%) single disulfide, 51.1% (±5.7%) double disulfide, and 5.1% (±0.3%) monomer.

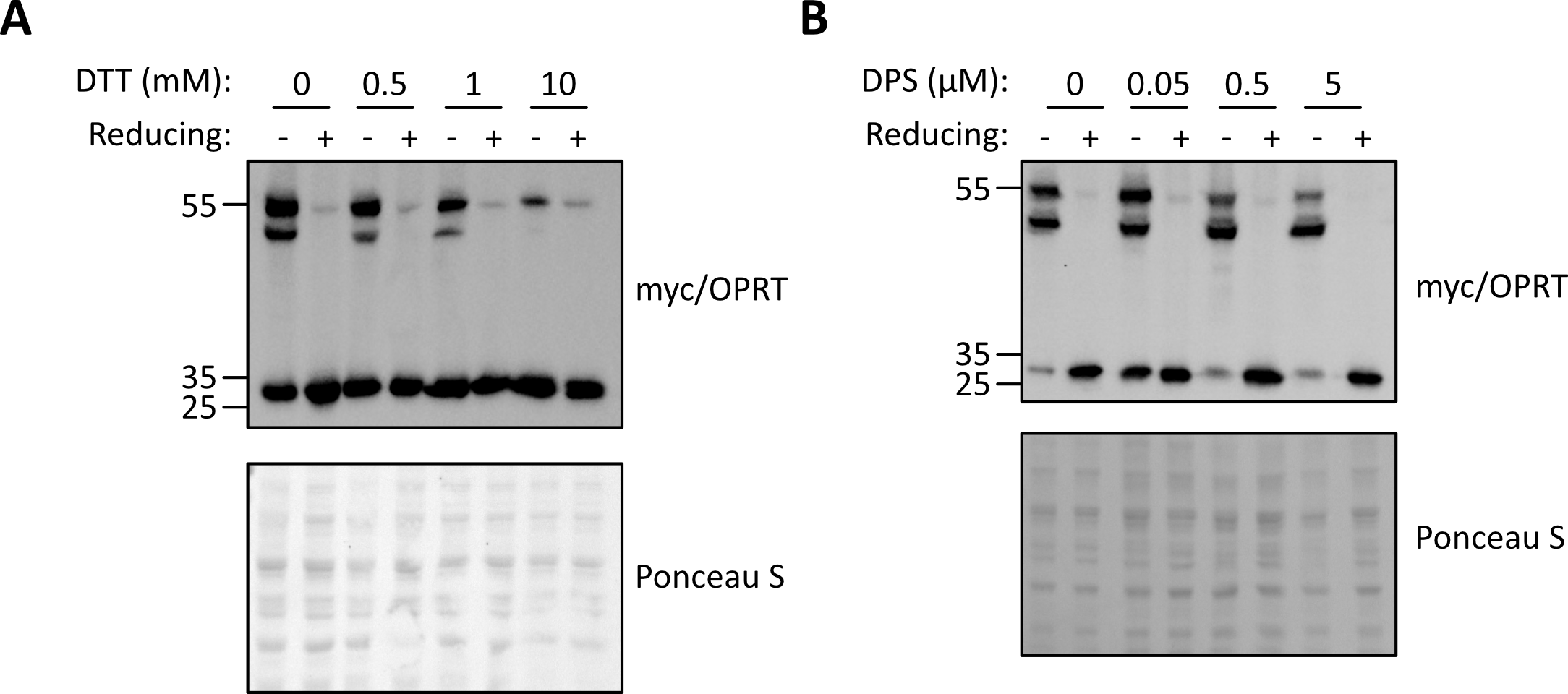

**Figure S6. Modulation of disulfide bonds using redox reagents.** (A) Western blot of yeast cells treated with the indicated concentrations of DTT before protein extraction. ‘Reducing’ indicates whether DTT was added (+) or omitted (-) in the sample buffer following extraction. (B) Western blot of yeast cells treated with the indicated concentrations of DPS before protein extraction. ‘Reducing’ indicates the same as in (A).

**Table S1. Primers used for mutagenesis**

| **Variant** | **Primer sequence** |
| --- | --- |
| R44C | 5'-GCTGTTTAATACCGGGTGCGATCTGGCACTGTT  5'-AACAGTGCCAGATCGCACCCGGTATTAAACAGC |
| D92C | 5'-GGAGCATCACGACCTGTGCCTGCCGTACTGCTT  5'-AAAGCAGTACGGCAGGCACAGGTCGTGATGCTCC |
| C96A | 5'-ACCTGGACCTGCCGTACGCCTTTAACCGCAAAGAAG  5'-CTTCTTTGCGGTTAAAGGCGTACGGCAGGTCCAGGT |
| C176A | 5'-GAAGTTGAGCGTGATTACAACGCCAAAGTGATCTCTATCATCAC  5'-GTGATGATAGAGATCACTTTGGCGTTGTAATCACGCTCAACTTC |
| R99G/K100S/K103S/H105G | 5'-TACTGCTTTAACGGCAGCGAAGCAAGCGACGGCGGTGAAGGCGGC  5'-GCCGCCTTCACCGCCGTCGCTTGCTTCGCTGCCGTTAAAGCAGTA |
| D124/125N | 5'-CGCGTAATGCTGGTAAATAATGTGATCACCGCCG  5'-CGGCGGTGATCACATTATTTACCAGCATTACGCG |

The primers used to create the variant described in the present paper are indicated (underlined are mutated positions).
